# Reading Profiles in Multi-site Data with Missingness

**DOI:** 10.1101/269555

**Authors:** Mark A. Eckert, Kenneth I. Vaden, Mulugeta Gebregziabher, Dyslexia Data Consortium

## Abstract

Children with reading disability exhibit varied deficits in reading and cognitive abilities that contribute to their reading comprehension problems. Some children exhibit primary deficits in phonological processing, while others can exhibit deficits in oral language and executive functions that affect comprehension. This behavioral heterogeneity is problematic when missing data prevent the characterization of different reading profiles, which often occurs in retrospective data sharing initiatives without coordinated data collection. Here we show that reading profiles can be reliably identified based on Random Forest classification of incomplete behavioral datasets, after the missForest method is used to multiply impute missing values. Results from simulation analyses showed that reading profiles could be accurately classified across degrees of missingness (e.g., ~5% classification error for 30% missingness across the sample). The application of missForest to a real multi-site dataset (n = 924) showed that reading disability profiles significantly and consistently differed in reading and cognitive abilities for cases with and without missing data. The results of validation analyses indicated that the reading profiles (cases with and without missing data) exhibited significant differences for an independent set of behavioral variables that were not used to classify reading profiles. Together, the results show how multiple imputation can be applied to the classification of cases with missing data and can increase the integrity of results from multi-site open access datasets.

## Reading Profiles in Multi-site Data with Missingness

Not all reading disabilities are the same. Children with reading disability can exhibit: 1) relatively specific phonological and spelling impairments (Fletcher, 2009; Lauterbach et al., 2017; Stanovich, 1988); 2) general language or language-learning impairments (Berninger & May, 2011; Bishop & Snowling, 2004), and 3) comprehension-specific impairments (Catts et al., 2012). These observations are generally consistent with hypotheses that phonological decoding and non-phonological language skills are unique and additive predictors of reading (Gough & Tunmer, 1986; Bishop & Snowling, 2004), although executive functions also appear to contribute to comprehension (Cain et al., 2004; Cutting et al., 2009; Locascio et al., 2010). The inclusion of children with these different reading disability profiles can be problematic for the outcomes of a study without a way to identify and account for reading different profiles that may have different etiologies (Samuelsson et al., 2007; Naples et al., 2009). For example, the study of retrospective multi-site data can include children with different reading profiles, particularly if sampling or ascertainment criteria were different across studies. This problem can be compounded if there is completely missing data from contributing research sites that could help differentiate reading disability profiles. Here we examined methods for dealing with missing data to identify reading profiles in a retrospective multi-site dataset.

Reading disability profiles have been characterized based on a variety of methods and measures that are sensitive to differences in reading skill, memory, perceptual, and general cognitive function (King et al., 2007; Miciak et al., 2015; Korkman & Häkkinen-Rihu, 1994; Kornilov & Grigorenko, 2017; Talcott et al., 2013). For example, trajectory analyses were used to differentiate reading disability cases with and without quite low general cognitive function (Kuppen & Goswami, 2016). Torpa et al. (2007) identified unique reading disability profiles, which included cases with and without low general cognitive function, using a latent class modeling approach. Or for example, Eckert et al., (2017) used a machine learning approach to verify clinician ratings of reading disability profiles that included relatively specific deficits in phonological decoding (Poor Decoder), low performance across reading and cognitive measures (Generally Poor Readers), and children with comprehension-specific reading disability (Poor Comprehenders).

While these reading disability profiles can be reliably classified on the basis of standardized test scores, missing data presents a major challenge for classification. This is particularly true for multi-site retrospective data analyses, when data collection was not coordinated before conducting studies. For example, missing a critical variable that is key for distinguishing between two profiles could prevent classification unless there is a method to deal with the missingness. On the other hand, a complete case analysis approach that ignores cases with missing scores could result in the exclusion of large numbers of children with a specific reading disability profile from multi-site datasets (e.g., Eckert et al., 2017). Such a strict inclusion policy introduces a false negative bias because of low power and this could be potentially mitigated by using multiply imputed data. The feasibility of identifying distinct reading profiles when there is missing data has not been examined, however.

In the current study, we examined three strategies for imputing missing data (mean replacement, multiple imputation with Predictive Mean Matching, multiple imputation with the missForest method) and the degree to which imputed data could then be used to reliably classify reading profiles. Multiple imputation is a statistically principled technique to estimate predictably missing values, usually by means of an informed regression model with an error term (Rubin, 1987; Little, 1988; Little & Rubin, 2002). The imputation model is used to produce multiple *(m)* simulated versions of the complete dataset that fill in missing values with new random draws that approximate the variance of the originally observed (i.e., non-missing) values. Each of the imputed datasets are separately analyzed and then the results across datasets are pooled (Little & Rubin, 2002). Multiple imputation produces low false negative and false positive rates when less than 30% of the data are missing across a sample (Vaden et al., 2012). Based on the intuition that a multivariate imputation model could be more effective at approximating multifactor patterns that are critical for classification analyses, we also tested the missForest algorithm for imputing missing data. The missForest method uses a Random Forest imputation model to estimate missing values based on decision trees instead of univariate regression models (Stekhoven & Buhlmann, 2012).

We examined the extent to which Poor Decoders, Generally Poor Readers, Poor Comprehenders, and Controls were consistently classified based on a retrospective multi-site dataset with missing data (Eckert et al., 2016; Eckert et al., 2017). We first examined classification accuracy in a synthetic dataset to demonstrate the relative advantage of missForest with the current dataset compared to other imputation approaches. We then applied missForest to a dataset with real missingness and compared the behavioral profiles of cases with and without missingness. Finally, we examined the extent to which the reading disability profiles could be validated using a set of independent measures of spelling and spatial cognition from the multi-site dataset.

## Methods

The data used in the current study were collected as part of a research project to develop methods for retrospective multi-site datasets that are increasingly a focus of data sharing initiatives. These data were contributed by members of the Dyslexia Data Consortium and included data from studies of reading disability group differences and dimensional studies of reading development. These data were contributed by members of the Dyslexia Data Consortium and included data from studies of reading disability group differences and dimensional studies of reading development. Specifically, these were studies in which participants were recruited for neuroimaging studies of dyslexia because they had received a clinical diagnosis of dyslexia, were enrolled in schools specializing in the education of children with dyslexia, or they were recruited from the local community for studies on reading development. Thus, the Dyslexia Data Consortium dataset is composed of participants who were recruited with varied sampling approaches. Additional information about the Dyslexia Data Consortium, including how to request the data and how to access the code used in this study can be found at www.dyslexiadata.org. Institutional Review Board (IRB) approval was obtained at each contributing site to share de-identified data. In addition, Medical University of South Carolina IRB approval was obtained to receive and analyze the de-identified data. We specifically requested word and pseudoword reading accuracy, rapid naming, and Verbal Comprehension (Verbal IQ) to determine the extent to which unique reading disability profiles could be reliably identified with these data based on the theoretical framework described above. We also requested any additional data that investigators could share. This resulted in a predominance of word reading accuracy and Verbal IQ measures, and other behavioral measures that were used in this study to provide external validity for the reading profiles.

The current study consisted of simulation and real dataset analyses. The simulation analyses were performed based on data that included 198 children from 8 different research sites (78 female; ages 6.39-12.85) without any missingness. These data were used in Eckert et al. (2017) to define reading profiles and allowed for the creation of simulated missingness to compare missingness approaches. Secondly, Random Forest was used to classify reading profiles in a sample of 726 children from 11 different research sites (334 female; ages 6.02-12.94 years), which excluded any cases from the Random Forest training dataset. Importantly, this dataset included cases with missing data that were imputed using the best method from the simulation results (missForest). The final analysis examined differences for behavioral measures that were not used to make the reading profile classifications from the 924 children whose data were included in the training and test datasets (11 research sites; 412 female; ages 6.02-12.94 years). Although there was extensive missingness in the combined validation dataset, this provided supporting evidence for the distinct reading profiles that were classified with observed and imputed data.

### Reading profile labels

The method for classifying reading profiles was developed in Eckert et al. (2017). These methods are presented here to explain how the reading profile labels were obtained and the accuracy of this approach. The Random Forest reading profiles classifications from Eckert et al. (2017) were used in the simulated and real data analyses for this study.

An expert-labeled training dataset was required to generate synthetic datasets and train a machine learning algorithm to classify reading profiles. The 198 children included in this training dataset had standardized scores from the (1) Woodcock-Johnson IIIR or Woodcock Reading Mastery Tests [Word Attack (pseudoword ID), Word Identification (real word ID), and Passage Comprehension; Woodcock, 1987; Woodcock et al., 2001], (2) Wechsler Intelligence Scales for Children or Wechsler Abbreviated Scales of Intelligence (Verbal Comprehension and Perceptual Reasoning or Verbal IQ; Wechsler, 1999, 2004), and (3) a standardized measure of Rapid Naming of letters or numbers (Wagner et al., 1999; Wolf & Denckla, 2005). Guided by the clinical and research expertise of a Neuropsychologist, the extant literature described above, and the age of the children, two clinically certified Speech-Language Pathologists labeled cases into the following theoretically defined profiles: 1) Poor Decoders, 2) Poor Comprehenders, 3) Generally Poor Readers, and 4) Controls. The resultant training dataset was composed of 58 Poor Decoders, 27 Poor Comprehenders, 14 Generally Poor Readers, and 99 Controls.

Random Forest was used to classify reading disability cases and controls, which matched the expert labels with 94% accuracy, based on leave-one-out cross validation tests (as reported in Eckert et al., 2017). The R statistics software (R version 3.3.1 with packages: caret v6.0.71, randomForest v4.6.12) was used to train the Random Forest classifier (parameters: *mtry* = 2; *number of trees* = 500) after a cross-validated tuning was performed (5-folds; 10 repeats) with the training dataset described above. The same classification training procedure was used for the synthetic and real data analyses in the current study.

### Simulation Analyses

#### Data synthesis

A simulation study was performed to evaluate classification accuracy when imputation was used to deal with missingness across different levels of missingness. The data synthesis tool *synthpop* R-package (v1.3.1; Nowok, Raab, Dibben, 2016) used in this study was originally developed to study epidemiologic, demographic, and health-related datasets without risk of re-identification as no case from an original dataset exactly re-occurs in the synthesized datasets. *synthpop* generates new datasets by estimating the value for each variable based on its association with the other variables in the data set, thus preserving the covariance structure across variables. We generated 1,000 synthetic datasets based on the expert-labeled training dataset using the default *synthpop* settings. These datasets had *N* = 500 participants and an equal number of labeled cases for each reading profile (Figure 1). Because expert labels for the reading profiles were propagated from the training dataset into the synthetic dataset, the accuracy of reading profile classification with the synthetic data was calculated by comparing the synthesized expert labels to the Random Forest classifications. Furthermore, the synthetic dataset allowed us to systematically remove known values to simulate missingness and assess the performance of different imputation methods.

**Figure 1.**
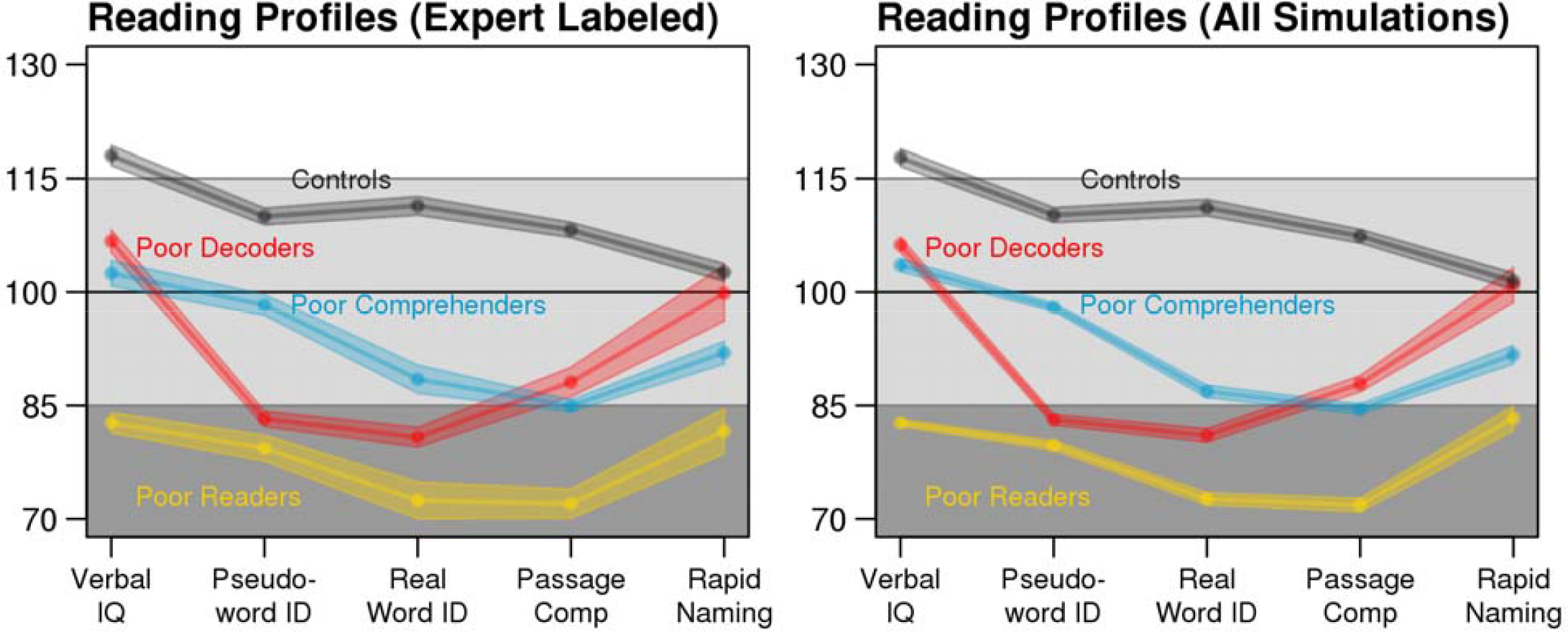
Consistent expert-labeled (left: n = 198) and synthetic data reading profiles (right: n = 500). Each point connected by a solid line is the mean test score and the shaded area depicts the standard error of the mean (SEM). For the synthetic data profiles, average test scores and SEM were calculated for each dataset with no missing data, then averaged across the 1,000 datasets.

#### Simulated missingness

To provide guidance about how the extent of missingness interacts with imputation performance and classification accuracy, we tested six levels of percent missingness within each synthetic dataset: 0, 10%, 20%, 30%, 40%, 50%. Imputation methods are most appropriate when used to estimate values that are missing predictably in relation to known participant characteristics (missing at random), or when those values are missing completely at random. Imputation methods are less appropriate when missingness is not at random, which occurs when the values are systematically related to the missingness, because the true range of values is greater than what is represented in the dataset. An example of missingness not at random would be missing comprehension scores because children had such severe reading disability that they could not perform the task. Here we focused on missingness at random, which can be a significant limitation in retrospective multi-site datasets.

Three variables in each simulation had missing data, which could include any variable that was used for profile classification (i.e., each participant’s age and five test scores). A missingness indicator (present = 0, missing = 1) represented the real or missing value and was used to ensure that each participant was missing no more than three values. Logistic regression was used to verify that sex and research site predicted missingness (*p* < 0.01). These demographic variables were selected to ensure missingness was predictable to evaluate these methods for missingness at random. Sex and age were selected, in particular, because they could explain why there would missing data for studies that are disproportionately represented by one sex or across age and therefore provide results that are generalizable to other studies.

#### Comparing imputation methods

Three imputation methods were tested to determine the approach that resulted in the highest reading profile classification accuracy when imputed datasets were submitted to a Random Forest classifier. (1) First, mean replacement was used to substitute each missing test score with the average observed score for that test. Although mean replacement is a common approach for analyzing data with missingness, this method is known to inflate Type I error rates by failing to approximate the distribution of values and underestimating its variance (Schafer & Graham, 2002). (2) Next, multiple imputation with Predictive Mean Matching (Little, 1988) was tested, which imputes missing values by drawing test scores from similar cases with no missing scores (R-package: MICE, v2.30; Van Buuren & Groothuis-Oudshoorn, 2011). (3) Finally, multiple imputation via the missForest method (R-package: missForest, v1.4; Stekhoven & Bühlmann, 2011) was tested. missForest fits an Random Forest model (Breiman, 2001) on all of the observed data to predict missing values for each factor with missing scores. Both Predictive Mean Matching and missForest imputation methods were used to generate ten imputed datasets that were each classified into reading profiles using a trained Random Forest model. Next, statistics were calculated for each of the *m* = 10 imputed datasets were pooled to form point and variance estimates (Little & Rubin, 2002). No such averaging was performed with mean replacement, because this approach can only generate one dataset. Critically, the synthetic reading profile labels were concealed from each of the imputation methods, so imputed test scores were informed by covariance in the test scores, participant age, participant sex, and research site, but not the reading profile labels.

Efficiency and classification accuracy were used to assess the imputation methods. Efficiency was calculated as the variance in the imputed scores divided by the variance in the scores from each test. Imputation methods that produce values that are more variable than the original data tend to produce false negative biases in univariate tests, while methods that reduce variance relative to the true values can produce false positive biases. Because we removed known values to simulate missingness, variance in the imputed test scores could be directly compared to the variance for the original test scores. The reading profile classification accuracy, sensitivity, and specificity were computed by comparing the Random Forest classification results to the reading profile labels in each synthetic dataset.

After determining that missForest resulted in the most accurate reading profile classifications, we assessed how missingness within each variable affected the classification accuracy for each reading profile when values were imputed with missForest. This involved selectively pooling profile-specific classification accuracy from each simulation that was missing for a specific variable. This allowed us to determine if each reading profile could be classified with confidence when a key variable that differentiated two profiles was missing. Each of these results was averaged across 1,000 simulations to produce reliable estimates for the effects of the amount of missingness and imputation methods on reading profile classification.

### Real Data Analyses

Multiple imputation was performed with the real dataset using missForest based on the simulation results described below. As noted above, independent classification tests were performed using a sample that shared no cases with the training dataset. These independent real data included 726 participants with missing values for Word Attack (0.4%), Word ID (0.1%), Passage Comprehension (16.1%), Verbal IQ (16.6%), and RAN (35.5%). Older children were more likely to be missing Passage Comprehension (age: *r*(724) = 0.28, *p* = 2.75E-14) and Verbal IQ data (age: *r*(724) = 0.19, *p* = 3.63E-7), while younger children were more likely to be missing RAN (age: *r*(724) = -0.07, *p* = 0.04). Children with higher Verbal IQ scores were more likely to be missing Passage Comprehension data (*r*(603) = 0.17, *p* =4.23E-5). In addition, research site significantly predicted missingness for Passage Comprehension (*Nagelkerke R-square* = 0.76, *p* = 1.90E-85), Verbal IQ (*Nagelkerke R-square* = 0.74, *p* = 1.96E-83), and RAN (*Nagelkerke R-square* = 0.55, *p* = 2.80E-74). Because the missingness largely resulted from research site differences in assessment plan, the missingness was predictable and missing at random. We had no information to suggest that missing data were missing not at random.

To validate reading profiles with independent behavioral measures, participants in the Random Forest training data were combined with the participants in the test dataset. This combined validation dataset included 924 participants and with the exception of Performance IQ scores (23.3% missing; Wechsler, 1999, 2004), there was substantial missingness for each variable used to independently validate the reading profiles. Specifically, Woodcock-Johnson Spelling (47.3% missing; Woodcock et al., 2001), Comprehensive Test of Phonological Processing Elision (39.8% missing; Wagner, Torgesen, & Rashotte, 1999), and Wechsler Digit Span (70.7% missing; Wechsler, 1999, 2004) measures exhibited too much missingness to reliably impute data. Caution is warranted in interpreting results from these data, but they were useful in illustrating distinct patterns of performance across the reading profiles.

## Results

### Synthetic Data Results

#### Imputation Diagnostics

Each of the expert-labeled reading profiles from the original dataset were well-approximated by the synthetic datasets used in the simulations (Figure 1, above). We compared the efficiency of each imputation method by calculating the ratio of the variance for the imputed test scores to the variance of the original test scores. Multiple imputation by Predictive Mean Matching most closely approximated the variance of the original dataset across levels of missingness (Figure 2). In contrast, Mean Replacement imputation demonstrated a steep decrease in efficiency as a function of percent missingness, simply because this method substituted all missing values with the mean observed value (variance = 0). missForest also underestimated the true variance in the data, although to a lesser extent than Mean Replacement. While underestimating variance can inflate the false positive rate for univariate statistics, the classification results below indicated that the Random Forest multivariate classifications were not biased when using the missForest imputed data.

**Figure 2.**
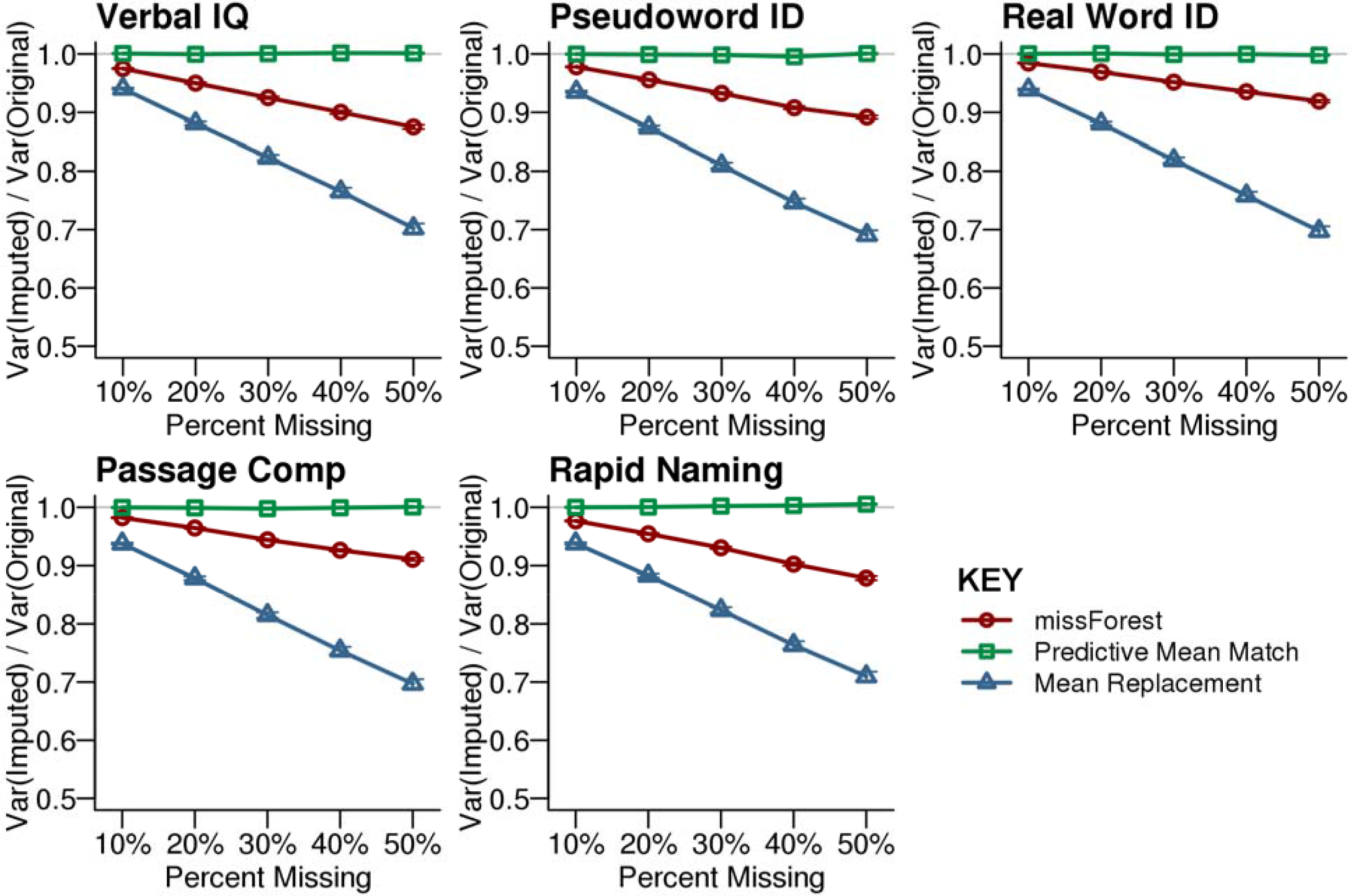
Efficiency for the imputed synthetic data. Efficiency is calculated by dividing the variance of the imputed dataset by the variance of the original dataset that had no missing data. Predictive Mean Matching provided the best approximation of the original variance across the tested levels of simulated missingness at random. The missForest imputed dataset exhibited less variance than the original dataset. Mean Replacement demonstrated the lowest efficiency with increasing missingness. Each point shows the average with SEM error bars calculated across 1,000 simulations.

#### Classification with simulated missingness

After simulated missing test scores were imputed with each method, each synthetic case was classified into a reading profile using RF. The highest classification accuracy was observed for the missForest imputed data, relative to the other missingness approaches. Across levels of simulated missingness, the smallest declines in classification accuracy, sensitivity, and specificity were observed for the missForest imputed data compared to the original synthetic data (Figure 3). These results suggest that classifications with missForest imputed data were the least biased and that missForest was most sensitive to the multivariate patterns that were present in the observed data.

**Figure 3.**
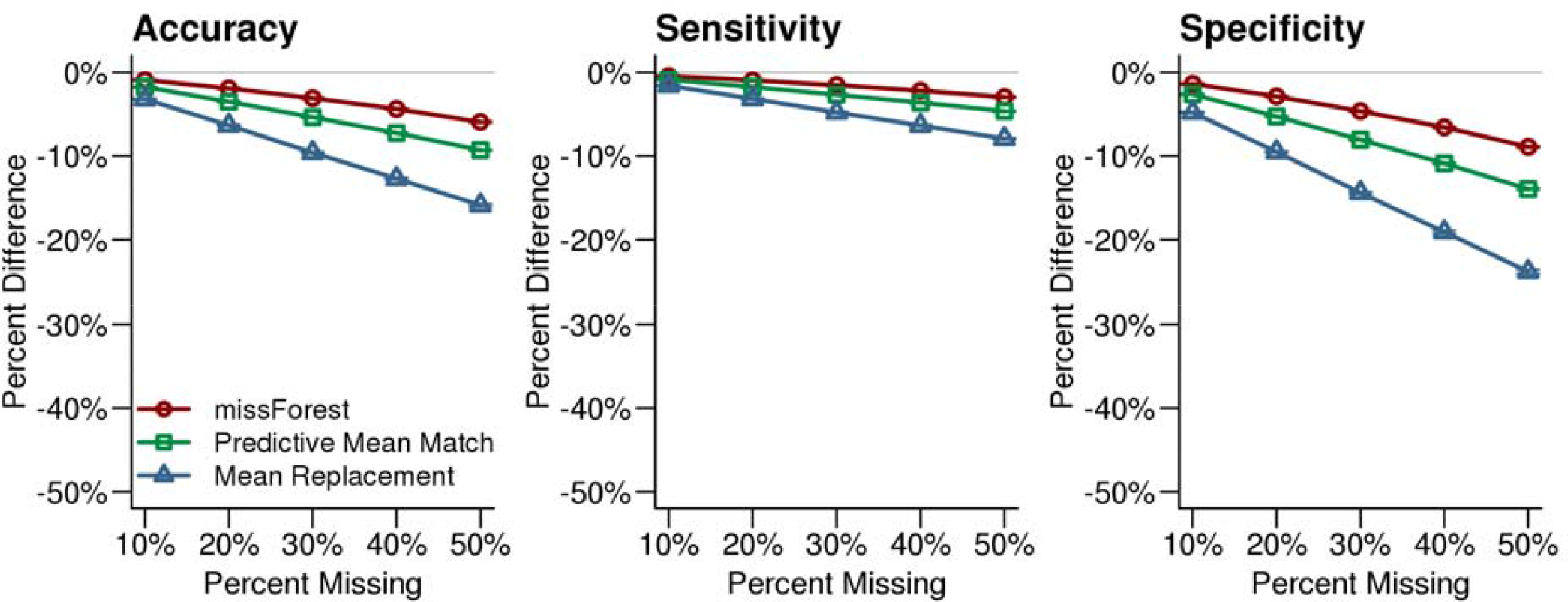
Reading profile classification accuracy for the imputed synthetic datasets. Reading profile classifications based on synthetic datasets with no missing data were compared to classifications that were performed after imputing missing values with missForest, Predictive Mean Matching, and Mean Replacement. The missForest imputation (red line) resulted in the most accurate classifications. The x-axis in each plot indicates the simulated percent missingness, while the y-axis indicates the reduction in classification accuracy, sensitivity, and specificity for the imputed datasets compared with the complete synthetic dataset. Each point shows the average with SEM error bars calculated over 1,000 simulations.

After determining that test scores imputed with missForest resulted in the highest classification accuracy, we examined classification accuracy for each reading profile when specific test scores were imputed using missForest (Figure 4). Poor Decoders and Poor Comprehenders were less accurately classified when pseudoword ID scores were among the missing data, while Generally Poor Readers were less accurately classified when Verbal IQ was missing. There was no single variable that negatively affected Control classification accuracy, which was high even for 50% missingness. Figure 4 shows that 30% missingness would increase classification error by ~5% across measures and reading profiles.

**Figure 4.**
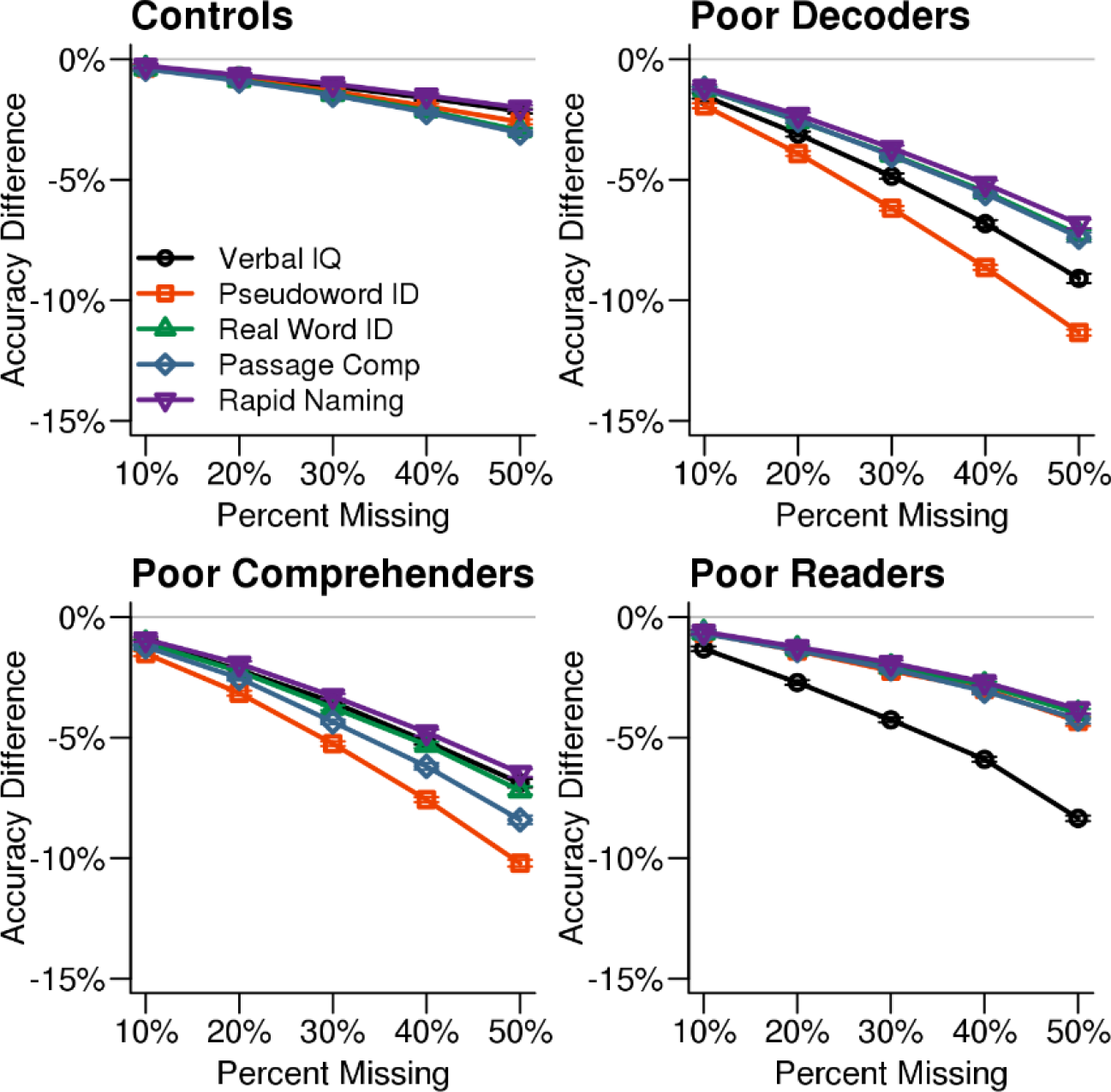
Classification accuracy by reading profiles and missing variables. Classification accuracy for each reading profile decreased when missForest was used to impute values for the measure that differentiates a reading profile from the others. For example, classification accuracy decreased more when Verbal IQ was missing for the Generally Poor Readers compared to the other test scores. The x-axis in each plot indicates the simulated percent missingness in the synthetic dataset, while the y-axis indicates the reduction in classification accuracy for the imputed datasets were compared with the original synthetic data with no missingness. Each point shows the average with SEM error bars calculated over 1,000 simulations.

### Real Data Results

#### Classification of real data with missingness

Multiple imputation increased the available sample size from 327 participants with complete data to 726 participants (Table 1), excluding the 198 cases used to train the classifier. The reading profiles exhibited similar patterns of behavioral strengths and weaknesses when comparing the training dataset to independent test dataset with no missing data and the test dataset with imputed scores (Figure 5), as well as the synthetic data (Figure 1). For example, Poor Decoder and Generally Poor Reader differences in Verbal IQ were significantly different for the cases whose data were imputed (ACA test dataset: *t*_(49)_ = 8.41, Cohen’s *d* = 2.54, *p* = 4.60E-11; MI: *t*(118.2) = 11.66, Cohen’s *d* = 2.58, *p* = 2.28E-21). These results show that missForest appear to have accurately estimated the values for missing variables that were critical for distinguishing the reading profiles (Figure 5).

Another demonstration of the effectiveness of missForest was the consistent reading profile classifications across the ten imputed datasets, as reflected in the low standard deviations for the classification labels (< 2 in Table 1). In addition, most of the correlations between behavioral measures were consistent across ACA and MI datasets (Supplemental Figure 1). For example, Verbal IQ and real word ID were significantly and positively related in Controls (ACA: *r*(232) = 0.55, *p* = 3.76E-20; MI: *r*(526.2) = 0.40, *p* = 3.69E-22), significantly and negatively related in Poor Comprehenders (ACA: r_(41)_ = -0.35, *p* = 0.02; MI: *r*(75.6) = -0.33, *p* = 0.004), and non-significantly related in Poor Decoders (ACA: *r*_(33)_ = -0.22, *p* = 0.21; MI: *r*(92.0) = 0.16, *p* = 0.13) and Generally Poor Readers (ACA: *r*_(14)_ = -0.45, *p* = 0.08; MI: *r*(24.2) = -0.36, *p* = 0.07). These results show that missForest preserved the covariance structure between the behavioral variables within each reading profile.

**Table 1.** Participant counts for each profile based on multiply imputed test scores or observed scores.

| Analysis Approach | Poor Decoders | Poor Comprehenders | Generally Poor Readers | Controls | Total N |
| --- | --- | --- | --- | --- | --- |
| Training Dataset | 58 | 27 | 14 | 99 | 198 |
| $\wedge$MI (mean ± sd) | 94 ± 1.1 | 77.6 ± 1.0 | 26.2 ± 0.8 | 528.2 ± 0.9 | 726 |
| ACA | 35 | 43 | 16 | 233 | 327 |

$\wedge$ MI: multiple imputation: Random Forest classifications based on missForest imputed data (pooled across ten imputed dataset). ACA: available case analysis. The independent test dataset for the MI and ACA analyses did not include participants from the reading profile training dataset.

**Figure 5.**
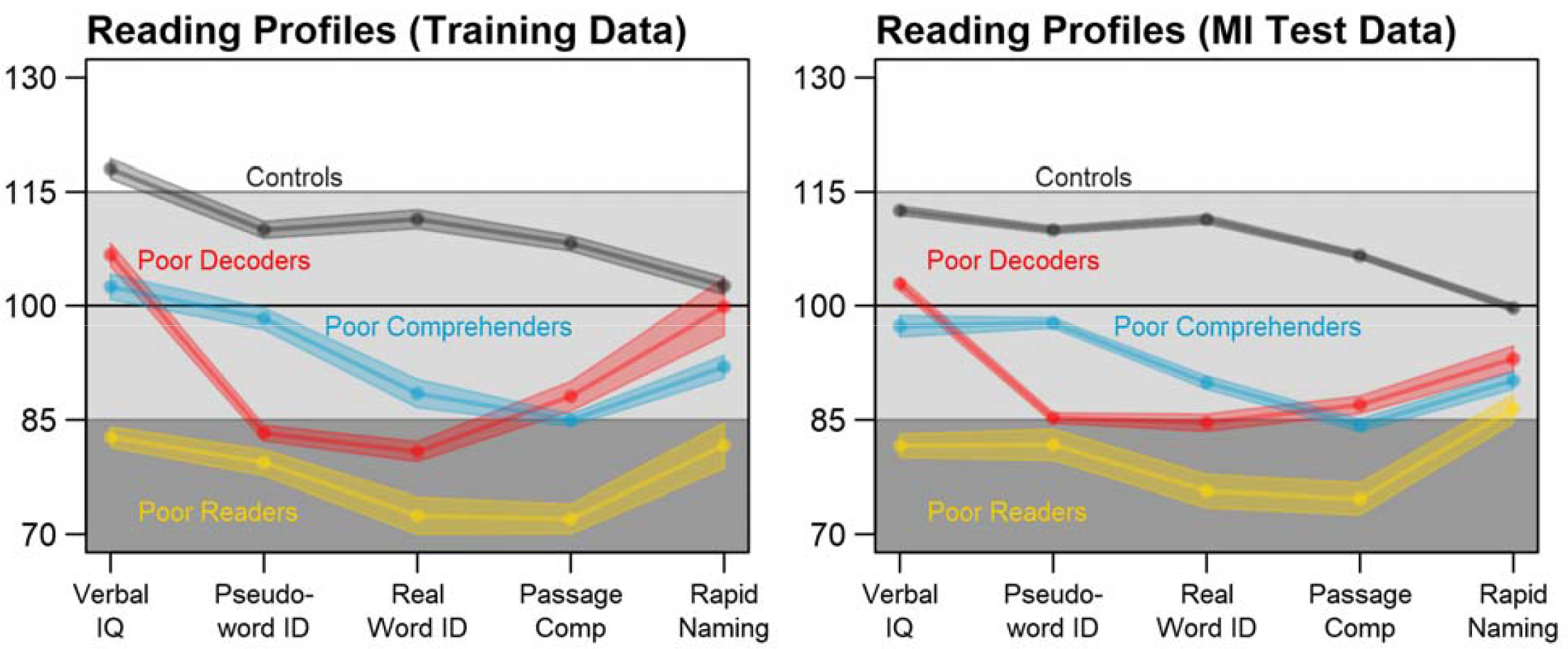
The same reading profiles are observed in the training dataset (N = 198) and MI test dataset (N = 726; pooled estimates from the *m* = 10 missForest imputations). Shaded area depicts the SEM.

### Validation of Reading Profiles

Behavioral measures that were not used to classify reading profiles were also examined to validate the profiles and determine the extent to which consistent differences were observed across complete case and multiple imputation analysis approaches. Figure 6 and Table 2 show that there were significant and unique reading profile differences for Performance IQ, Spelling, Elision, and Digit Span measures. Reading disability cases exhibited consistently lower scores compared to controls across these measures. Generally Poor Readers were consistently the lowest performers across measures. Generally Poor Readers and Poor Decoders demonstrated significantly lower Elision performance compared to Poor Comprehenders. In addition, Poor Decoders could be differentiated from Generally Poor Readers because of significantly better Performance IQ and Digit Span performance than the Generally Poor Readers. These results were observed for both the complete case and multiply imputed datasets. Caution is appropriate in interpreting the patterns of reading profile differences for these variables because of the significant extent of missingness, which prevented these data from being imputed or informing the Random Forest classification model. Nonetheless, the results indicate that the four reading profiles could be distinguished with an independent set of behavioral measures, thereby providing validation of the four reading profiles in this multi-site dataset.

**Table 2.**
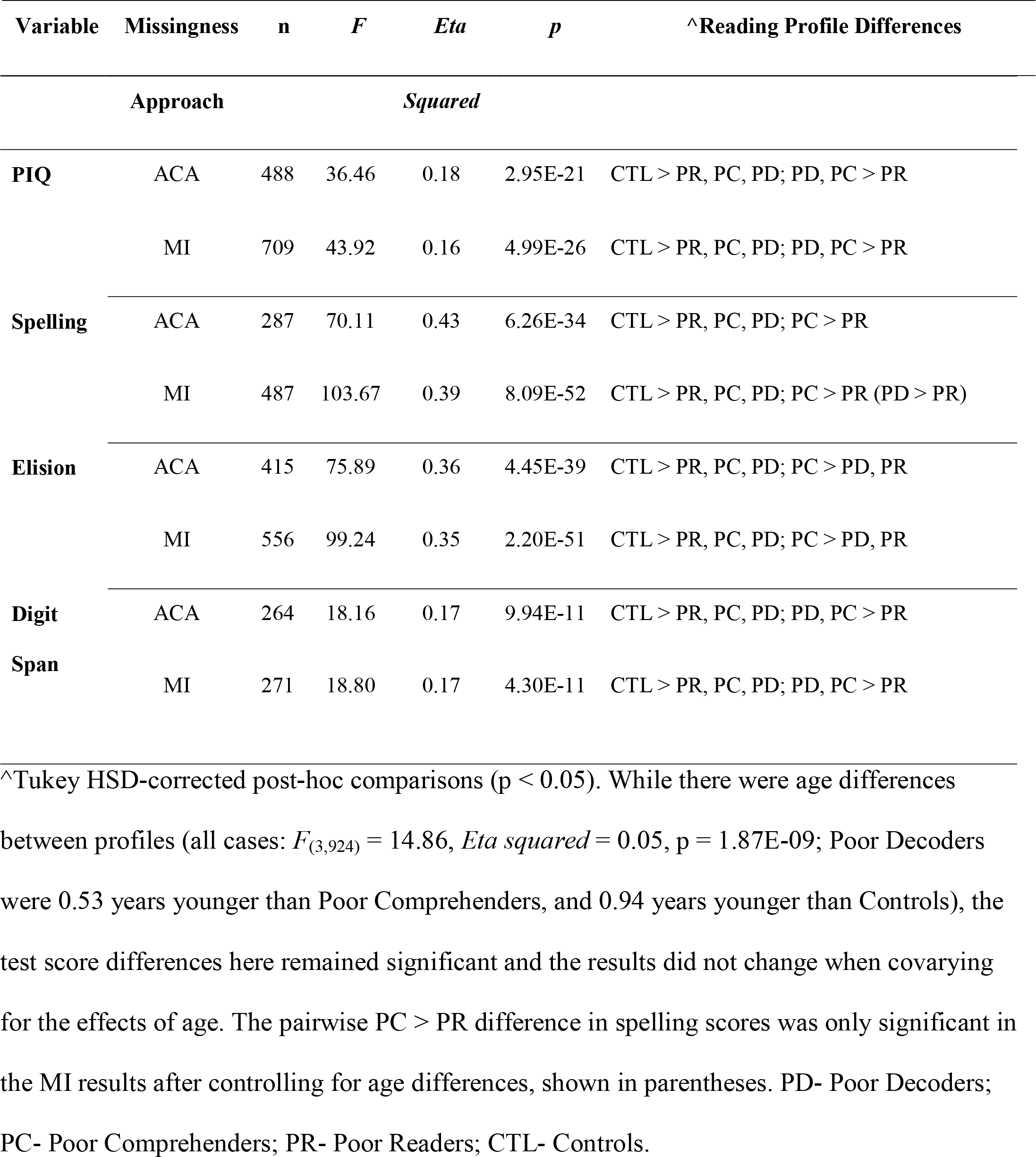
Reading profile differences were observed for complete case (ACA) and multiply imputed (MI) data for behavioral measures that were not used to classify the reading profiles.

**Figure 6.**
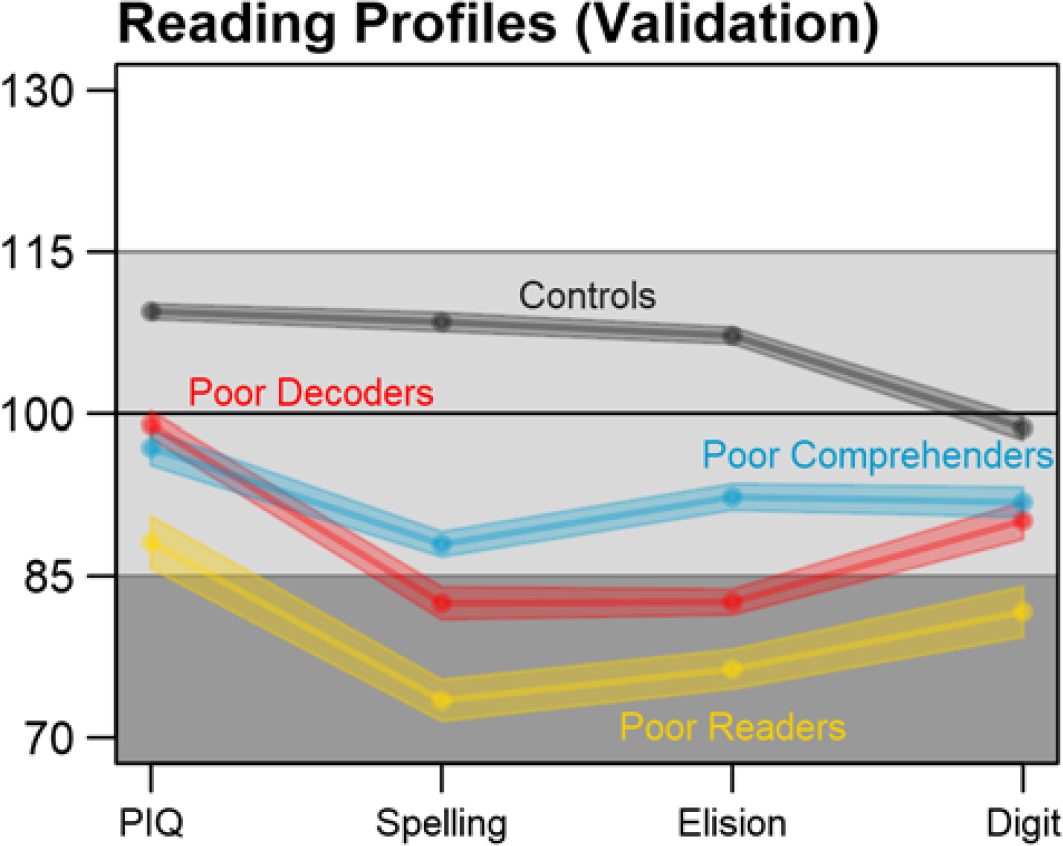
Reading profile validation with independent measures. The reading profiles, including cases with imputed data, could be differentiated using additional test scores that were available in the multi-site dataset, but that had too much missingness to be imputed and used for reading profile classification. Performance IQ = PIQ. Average test scores and SEM were calculated for each imputation, then pooled over the *m* = 10 imputed datasets.

## Discussion

Children with reading disability exhibit heterogeneous deficits in component reading skills that contribute to the presentation of unique reading profiles. These reading profiles appear to have importance for understanding the etiologies of reading disability because they have been linked to different patterns of brain morphology (Bailey et al., 2016; Jednoróg et al., 2014; Leonard et al., 2006; Leonard & Eckert, 2008) and genetic findings (Samuelsson et al., 2007; Naples et al., 2009). Although we were not able to identify the etiology of the reading profiles examined in this study, or characterize the presence of any additional reading profiles in the current dataset (Tamboer et al., 2016; Torppa et al., 2007; Wolff, 2010; Kornilov & Grigorenko, 2017), we have demonstrated that the reading profiles described in the current study can be reliably and consistently classified with ~30% missingness. Classification accuracy exhibited limited decline when missing data were imputed with missForest, even when key variables that differentiate reading profiles were missing. We also validated the reading profiles with behavioral data that were not used for classification. The methods used in this project could be leveraged to characterize or control for behavioral heterogeneity in reading studies, and can be applied in studies of other complex behavioral disorders.

### Simulation Results

The simulation results provided critical information about the degree to which imputed data could accurately characterize variance compared to the real data, and allowed for comparisons of reading profile classification accuracy between imputation methods. Predictive Mean Matching was the most accurate imputation method based on variance approximation of the original dataset across percent levels of missingness, which indicates that it should provide relatively limited false positive results in univariate statistic tests. Nonetheless, the data imputed using missForest resulted in the highest reading profile classification accuracy across levels of missingness. The missForest imputed data also resulted in less biased classifications based on higher sensitivity and higher specificity compared to the other two methods. In the context of multivariate analyses such as Random Forest classification, missForest appears to have advantages over the other imputation methods even though the variability of imputed values was reduced, which could increase false positive rates for univariate tests. Thus, Predictive Mean Matching may be a preferable multiple imputation approach for univariate analyses, while missForest appeared to better capture the relations between variables and was preferable for multivariate classification analyses.

Perhaps not surprisingly, reading profile misclassification increased when a missing variable was critical for differentiating one reading profile from the others. For example, Generally Poor Reader misclassification was more likely when Verbal IQ was missing, while Poor Comprehender and Poor Decoder misclassification were more likely when pseudoword ID was missing. However, there were only ~5% more classification errors with 30% simulated missingness for these critical variables. These results suggest that missForest can be applied to a real dataset to optimize reading profile classification accuracy.

### Real Dataset Results

Again, we sought to develop methods that could help identify homogeneous cases of reading disability in a retrospective dataset, despite missing data. There is some evidence that reading disability classifications using cut-points can be inconsistent depending on the types of measures, in part because of inconsistent measurement reliability (Miciak et al., 2015). Measurement reliability and standardization could affect the validity of imputation and classification. The measures used for classification in the current study are widely used in the reading disability community and have demonstrated good concurrent and predictive validity (Fuchs et al., 2004; Fuchs & Young, 2006; Schatschneider et al., 2004).

We used a multivariate approach to identify patterns of performance across component reading skills, which was guided by a theoretical framework (Bishop & Snowling, 2004), clinical expertise, and empirical evidence for unique reading profiles. We used measures that are commonly administered in reading disability studies (e.g., Woodcock-Johnson Word Attack) to leverage as much of the multi-site dataset as possible. One drawback to using only real word and pseudoword accuracy measures is the absence of reading efficiency, orthography, and expressive or receptive syntax measures to help guide the classifications. These measures could have enhanced multiple imputation and reading profile classifications.

While our results support the premise for characterizing three different reading disability groups, the profile plot figures suggest that combinations of continuous effects within cognitive domains produces these different profiles. For example, the profile plots of the Generally Poor Readers and Poor Comprehenders in Figures 1 and 5 are largely parallel. It is the magnitude of impairment that appears to differentiate these profiles. In addition, the pattern of associations between Verbal IQ and reading measures was similar consistent for the Generally Poor Readers and Poor Comprehenders (Supplemental Figure 1). These results could indicate that Poor Comprehenders and Generally Poor Readers have a common latent impairment that was not assessed, such as poor organization or planning skills, which affects multiple functions and has a more exaggerated effect for challenging tasks that require support from multiple neural systems (i.e., Passage Comprehension). This idea could be consistent with evidence that children exhibiting comprehension-specific reading disability also exhibit relatively low executive function performance (Locascio et al., 2010), which may explain this unexpectedly low oral language performance (Cutting et al., 2009). Low oral language performance (e.g., receptive syntax understanding) would also be expected in the Generally Poor Readers (Oral and Written Language Learning Disability; Berninger, 2008; Berninger & May, 2011). We predict that the magnitude of executive function impairment creates differences across multiple behavioral measures between Generally Poor Readers and Poor Comprehenders. The Generally Poor Readers were not classified into the Poor Comprehenders profile, in part, because they also exhibited behavioral similarity to the Poor Decoders, as we discuss below.

### Independent Validation

We leveraged data that were not used for imputation and reading profile classification to demonstrate that the reading profiles could be differentiated with Performance IQ, Elision, Spelling, and Digit Span measures. The same pattern of behavioral differences was observed for cases with complete data and for cases who were classified with imputed data. The only difference was the larger magnitude of *F* statistics for the imputed datasets, although not *eta squared* effect sizes, suggesting that the reading profile differences were stable.

The validation data and results were limited to the extent that there were too many cases with missing data for reliable imputation results. Nonetheless, these variables and results provided evidence of differences between the reading profiles that included: 1) Generally Poor Readers who had consistently low scores compared to Controls and lower Performance IQ and Digit Span compared to Poor Decoders and Poor Comprehenders; 2) Poor Decoders who had lower Elision than the Poor Comprehenders, but did not differ with Poor Comprehenders in Performance IQ and Digit Span performance; and 3) Poor Comprehenders who performed consistently lower than Controls, but as noted above, had better performance for the Spelling and Elision measures compared to the Generally Poor Readers and better Elision than the Poor Decoders.

In contrast to Figures 1 and 5 in which Generally Poor Readers and Poor Comprehenders had similar patterns and differed in magnitude of impairment, Figure 6 shows similar patterns for Generally Poor Readers and Poor Decoders that differed in the magnitude of impairment. In addition, the pattern of associations between reading measures was also consistent for the Generally Poor Readers and Poor Decoders (Supplemental Figure 1). These results suggest that impairments associated with Poor Decoders also contribute to the reading problems that Generally Poor Readers experience. Thus, Generally Poor Readers could experience executive function problems that affect Poor Comprehenders, as well as phonological processing problems that affect Poor Decoders. This interpretation is broadly consistent with the premise that Generally Poor Readers experience a combination of decoding and comprehension problems (Gough & Tunmer, 1986) or non-phonological language problems (Bishop & Snowling, 2004).

### Limitations

The missingness and classification methods used in this study were effective despite an unbalanced number of participants in each reading profile. Users of these methods should be cautious in their implementation with relatively small datasets when it is possible that the cases belonging to the under-sampled profile could be misclassified because of a relatively limited contribution of these cases to the imputation model. We also acknowledge that the three reading disability profiles examined in this study may not be a comprehensive characterization of different reading profiles. Again, we did not have access to reading efficiency or fluency measures that might have further differentiated the profiles. We were able to independently validate the three reading profiles, but also note again that this study was limited by the more extensive missingness for the variables used to validate our results.

## Conclusions and Future Study

Researchers are increasingly motivated to share data in open access databases. These data sharing and scientific transparency initiatives can advance the pace of discoveries and enhance scientific integrity, but retrospective analysis of data collected without coordination can be challenging for many reasons. Specifically, children with different reading disability profiles can be included in multi-site datasets, particularly if different ascertainment or sampling criteria were used between research sites. This sampling heterogeneity can introduce variance and challenge our ability to make inferences from multi-site results. Ideally, investigators could control for this heterogeneity, but this is difficult when there is missing data that would help differentiate reading profiles. The current study demonstrates that consistent patterns of reading disability can be identified in a multi-site retrospective dataset, even when there is missing data and different numbers of children with each reading profile.

We used simulations and real data to show that reading profiles can be classified with a high degree of accuracy when missing data are imputed using the missForest method. The sample size of the real dataset was almost doubled by adding participants with missing data to the participants with complete data. This provides a clear example of how the missForest imputation method could increase statistical power and reduce the false negative rate. Including cases with missing data also potentially increases the quality of inference because of more representative sampling, as demonstrated in Figure 5.

Additional work and perhaps an expanded set of behavioral measures that includes executive function, visual perception, and cognitive processing speed, as well as developmental and family histories that could provide additional validation for the reading profiles. We also emphasize that the methods used in this current study are not designed to classify a single participant for clinical reasons or for case studies. These methods should be used to enhance the validity and inference from group level statistics. An important and broader implication of the findings is that the methods demonstrated here could be applied to the study of other complex developmental and age-related disorders where there is evidence for the expression of unique profiles or phenotypes. Thus, we anticipate that the combined multiple imputation and classification approach described here can be applied to open access datasets that are composed of heterogeneous cases.

## References

Bailey, S., Hoeft, F., Aboud, K., & Cutting, L. (2016). Anomalous gray matter patterns in specific reading comprehension deficit are independent of dyslexia. Annals of Dyslexia, 66(3), 256-274.

Berninger, V. W. (2008). Defining and differentiating dysgraphia, dyslexia, and language learning disability within a working memory model. Brain, Behavior, and Learning in Language and Reading Disorders, 103-134.

Berninger, V. W., & May, M. O. (2011). Evidence-based diagnosis and treatment for specific learning disabilities involving impairments in written and/or oral language. Journal of Learning Disabilities, 44(2), 167-183.

Bishop, D. V., & Snowling, M. J. (2004). Developmental dyslexia and specific language impairment: Same or different? Psychological Bulletin, 130(6), 858-886.

Breiman, L. (2001). Random forests. Machine Learning, 45(1), 5-32.

Cain, K., Oakhill, J., & Bryant, P. (2004). Children’s reading comprehension ability: Concurrent prediction by working memory, verbal ability, and component skills. Journal of Educational Psychology, 96(1), 31.

Catts, H. W., Compton, D., Tomblin, J. B., & Bridges, M. S. (2012). Prevalence and nature of late-emerging poor readers. Journal of Educational Psychology, 104(1), 166-181.

Cutting, L. E., Materek, A., Cole, C. A., Levine, T. M., & Mahone, E. M. (2009). Effects of fluency, oral language, and executive function on reading comprehension performance. Annals of Dyslexia, 59(1), 34-54.

Eckert, M. A., Berninger, V. W., Vaden, K. I., Gebregziabher, M., & Tsu, L. (2016). Gray matter features of reading disability: A combined meta-analytic and direct analysis approach. ENeuro, 3(1).

Eckert, M. A., Vaden, K. I., Maxwell, A. B., Cute, S. L., Gebregziabher, M., & Berninger, V. W. (2017). Common brain structure findings across children with varied reading disability profiles. Scientific Reports, 7(1), 6009.

Fletcher, J. M., Lyon, G. R., Fuchs, L. S., & Barnes, M. A. Learning Disabilities: From Identification to Intervention. (Guilford Publications, 2006).

Fletcher, J. M. (2009). Dyslexia: The evolution of a scientific concept. Journal of the International Neuropsychological Society, 15(4), 501–508.

Fuchs, L. S., Fuchs, D., & Compton, D. L. (2004). Monitoring early reading development in first grade: Word identification fluency versus nonsense word fluency. Exceptional Children, 71(1), 7-21.

Fuchs, D., & Young, C. L. (2006). On the irrelevance of intelligence in predicting responsiveness to reading instruction. Exceptional Children, 73(1), 8-30.

Gough, P. B., & Tunmer, W. E. (1986). Decoding, reading, and reading disability. Remedial and Special Education, 7(1), 6-10.

Jednoróg, K., Gawron, N., Marchewka, A., Heim, S., & Grabowska, A. (2014). Cognitive subtypes of dyslexia are characterized by distinct patterns of grey matter volume. Brain Structure and Function, 219(5), 1697-1707.

King, B., Wood, C., & Faulkner, D. (2007). Sensitivity to auditory and visual stimuli during early reading development. Journal of Research in Reading, 30(4), 443-453.

Korkman, M., & Hakkinenrihu, P. (1994). A new classification of developmental language disorders (DLD). Brain and Language, 47(1), 96-116.

Kornilov, S. A., & Grigorenko, E. L. (2017). What reading disability? Evidence for multiple latent profiles of struggling readers in a large Russian sibpair sample with at least one sibling at risk for reading difficulties. Journal of Learning Disabilities: 0022219417718833

Kuppen, S. E., & Goswami, U. (2016). Developmental trajectories for children with dyslexia and low IQ poor readers. Developmental Psychology, 52(5), 717-734.

Lauterbach, A. A., Park, Y., & Lombardino, L. J. (2017). The roles of cognitive and language abilities in predicting decoding and reading comprehension: comparisons of dyslexia and specific **language** impairment. Annals of Dyslexia, 67(3), 201-218.

Leonard, C. M., & Eckert, M. A. (2008). Asymmetry and dyslexia. Developmental Neuropsychology, 33(6), 663-681.

Leonard, C. M., Eckert, M. A., Given, B., Berninger, V., & Eden, G. (2006). Individual differences in anatomy predict reading and oral language impairments in children. Brain, 129(12), 3329-3342.

Little, R. J. A. (1988). Missing data adjustments in large surveys. Journal of Business Economics and Statistics, 6, 287-301.

Little, R. J. A., & Rubin, D. B. (2002). Statistical Analysis with Missing Data. 2nd ed. Hoboken, NJ: Wiley Interscience.

Locascio, G., Mahone, E. M., Eason, S. H., & Cutting, L. E. (2010). Executive dysfunction among children with reading comprehension deficits. Journal of Learning Disabilities, 43(5), 441-454.

Miciak, J., Taylor, W. P., Denton, C. A., & Fletcher, J. M. (2015). The effect of achievement test selection on identification of learning disabilities within a patterns of strengths and weaknesses framework. School Psychology Quarterly, 30(3), 321.

Naples, A. J., Chang, J. T., Katz, L., & Grigorenko, E. L. (2009). Same or different? Insights into the etiology of phonological awareness and rapid naming. Biological Psychology, 80(2), 226-239.

Nowok, B., Raab, G. M., & Dibben, C. (2016). synthpop: Bespoke creation of synthetic data in R. Journal of Statistical Software, 74(11), 1-26.

Rubin, D. B. Multiple Imputation for Nonresponse in Surveys. (J. Wiley & Sons, New York, 1987).

Samuelsson, S., Olson, R., Wadsworth, S., Corley, R., DeFries, J. C., Willcutt, E., … Byrne, B. (2007). Genetic and environmental influences on prereading skills and early reading and spelling development in the United States, Australia, and Scandinavia. Reading and Writing, 20(1), 51-75.

Schafer, J. L., & Graham, J. W. (2002). Missing data: Our view of the state of the art. Psychological Methods, 7(2), 147–177.

Schatschneider, C., Fletcher, J. M., Francis, D. J., Carlson, C. D., & Foorman, B. R. (2004).Kindergarten prediction of reading skills: A longitudinal comparative analysis. Journal of Educational Psychology, 96(2), 265.

Stanovich, K. E. (1988). Explaining the differences between the dyslexic and the garden-variety poor reader: The phonological-core variable-difference model. Journal of Learning Disabilities, 21(10), 590-604.

Stekhoven, D. J., & Bühlmann, P. (2011). MissForest - non-parametric missing value imputation for mixed-type data. Bioinformatics, 28(1), 112-118.

Tamboer, P., Vorst, H. C., & Oort, F. J. (2016). Five describing factors of dyslexia. Journal of Learning Disabilities, 49(5), 466-483.

Torppa, M., Tolvanen, A., Poikkeus, A. M., Eklund, K., Lerkkanen, M. K., Leskinen, E., & Lyytinen, H. (2007). Reading development subtypes and their early characteristics. Annals of Dyslexia, 57(1), 3-32.

Vaden, K. I., Gebregziabher, M., Kuchinsky, S. E., & Eckert, M. A. (2012). Multiple imputation of missing fMRI data in whole brain analysis. Neuroimage, 60(3), 1843-1855.

Van Buuren, S., & Groothuis-Oudshoorn, K. (2011). mice: Multivariate Imputation by Chained Equations in R. Journal of Statistical Software, 45(3), 1-67.

Wagner, R. K., Torgesen, J. K., & Rashotte, C. A. Comprehensive Test of Phonological Processing (Pro-Ed Inc, 1999).

Wechsler, D. Wechsler Abbreviated Scale of Intelligence (The Psychological Corporation., 1999).

Wechsler, D. The Wechsler Intelligence Scale for Children - 4th edition. (Pearson Assessment, 2004).

Wolf, M., & Denckla, M. B. Rapid Automatized Naming and Rapid Alternating Stimulus Tests (RAN/RAS). (Pro-Ed Inc, 2005).

Wolff, U. (2010). Subgrouping of readers based on performance measures: A latent profile analysis. Reading and Writing, 23(2), 209-238.

Woodcock, R. W., Mather, N., McGrew, K. S., & Shrank, F. A. Woodcock-Johnson III Tests of Cognitive Abilities. (Riverside Publishing, 2001).

Woodcock, R. Woodcock Reading Mastery Test: revised. (American Guidance Service, 1987).

